# Wavelet coherence phases decode the universal switching mechanism of Ras GTPase superfamily

**DOI:** 10.1101/2020.08.15.252247

**Authors:** Zenia Motiwala, Anand S. Sandholu, Durba Sengupta, Kiran Kulkarni

## Abstract

Ras superfamily GTPases are molecular switches which regulate critical cellular processes. Extensive structural and computational studies on these G proteins have established a general framework for their switching mechanism, which involves conformational changes in their two loops, Switch I and Switch II, upon GTP binding and hydrolysis. Since the extent of these conformational changes is not uniform amongst the members of the Ras superfamily, there is no generic *modus operandi* defining their switching mechanism. Here, we have developed a novel approach employing wavelet coherence analysis to correlate the structural changes with their functional states. Our analysis shows that the structural coupling between the Switch I and Switch II regions is manifested in terms of conserved wavelet coherence phases, which could serve as useful parameters to define functional states of the GTPases. In oncogenic GTPases mutants, this phase coupling gets disentangled, which perhaps provides an alternative explanation for their aberrant function. We have tested the statistical significance of the observed phase angle correlations on multiple switch region conformers, generated through MD simulations.

## Introduction

Proteins utilize reversible transformations, such as covalent modifications (example: phosphorylation) and ligand induced conformational changes, to exert signal transduction events. For instance, Ras superfamily of GTPases act as biomolecular switches to regulates diverse biological functions such as cell migration, proliferation and fate. They achieve this switching mechanism by shuttling between GDP bound ‘OFF’ and GTP bound ‘ON’ states (Fig 1A). Comparative analysis of structures in different ligand bound states has proved to be a useful technique to gain functional insights. However, in few cases, simple structure comparison has proved to be inadequate to derive generalized inferences, as the observed differences between structures are either not quantifiable or they exhibit complex divergence patterns. This is demonstrated from the Ras superfamily of GTPases, which comprises of 5 families: Rho, Ras, Rab, Ran and Arf GTPase (Goitre *et al*, 2014; Valencia *et al*, 1991; Wennerberg *et al*, 2005a). All the members of this superfamily possess the conserved nucleotide binding G domain, containing a 6-stranded mixed β-sheet surrounded by five α-helices. The G-domain harbors five conserved sequence motifs termed G1-G5, which stabilize the nucleotide-protein interactions. Out of these motifs, the P loop GxxxxGKT (G1) and the two switch regions (SWI & SWII), containing the xT/Sx (G2) and DxxG (G3) motifs, interact with phosphates of the nucleotide to function as sensors for the presence of the gamma-phosphate (Goitre *et al*, 2014; Valencia *et al*, 1991; Wennerberg *et al*, 2005a). Structures of GDP and GTP bound proteins have shown that, primarily the SWI and SWII regions of the protein undergo conformational changes to deploy the conserved “loaded-spring” mechanism where release of the γ-phosphate after GTP hydrolysis allows the two switch regions to relax into the GDP specific conformation (Fig S1A). These conformational changes are attributed to loss of two hydrogen bonds between γ-phosphate oxygens and the main chain NH groups of the invariant Thr and Gly residues (Thr35/ Gly60 in Ras) in SWI and SWII, respectively (Fig S1B). Thus it was inferred that the functional states of Ras superfamily GTPases is the consequence of the conformational changes in their SWI and SWII regions (Pylypenko et al, 2018; Vetter & Wittinghofer, 2001).

**Figure 1.**
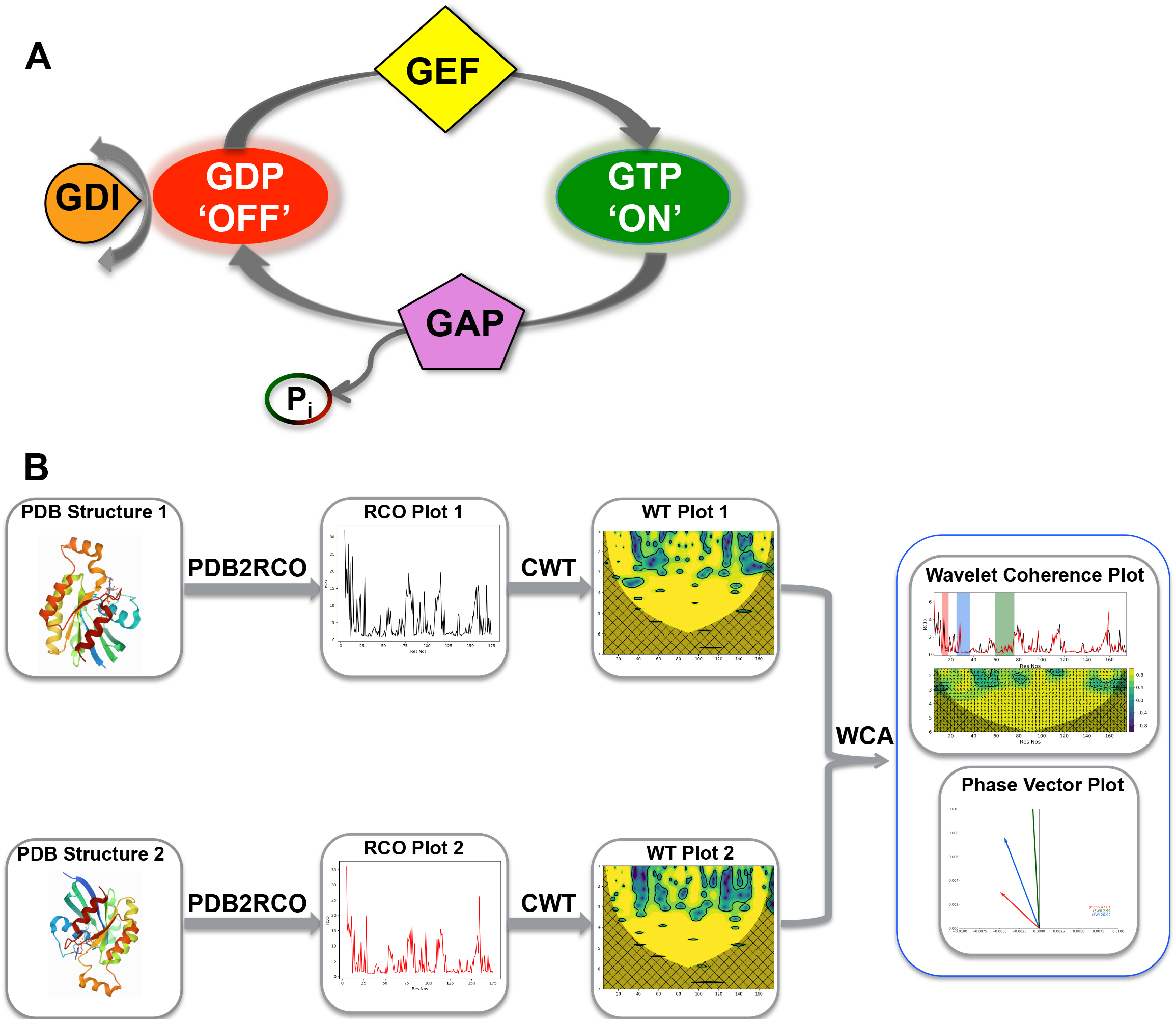
Schematic representation of GTPase switching and Wavelet based methodology to study this switching in the current study. **(A)** Switching mechanism of the Ras superfamily GTPases. The switching between GTP bound ON and the GDP bound OFF state is regulated by Guanine Nucleotide Exchange Factors (GEFs) and GTPase Activating Proteins (GAPs); GEFs catalyze the exchange of their bound GDP with GTP and GAPs promote intrinsic GTP hydrolysis. The Rho GTPase subfamily, has an additional level of regulation involving Guanine Dissociation Inhibitors (GDIs), which sequester GDP bound Rho GTPases in the cytosol. **(B)** Flowchart of the wavelet based methodology developed for analyzing the GTPases.

However, the quantum of conformational change in SWI and SWII regions, upon change in their bound nucleotides, vary across different sub-families. For example, amongst GTPases considered here, Rho, Ras, Rab and Arf, the Rho family exhibits insignificant changes in the conformation of SWI & SWII regions, whereas in the other three it is little more dramatic (Fig 2B, 2D, 2F and 2H). Therefore, correlating the functional states of GTPases from their SWI & SWII conformations is really challenging. This conundrum further poses difficulties in understanding mechanistics of GTPase interactions with their regulators and effectors. Furthermore, the classical way of ascertaining their functional states (ON & OFF) in terms of loss or gain of hydrogen bonds between γ-phosphate oxygens of the GDP/GTP (S1B) with the protein would be an insufficient parameter, as GTPases have been proposed to exhibit intermediate conformational states such as “ON like” state (Pálfy *et al*, 2019; Gorfe *et al*, 2008; Kapoor & Travesset, 2014; Li *et al*, 2018a; Muraoka *et al*, 2012; Anand *et al*, 2013; Prakash & Gorfe, 2013) This question becomes even more pertinent while deciphering the impact of oncogenic mutants on the functioning of GTPases, as the crystal structures of mutants at positions 12, 13 or 61 (numbering as per Cdc42 sequence), show no significant conformational deviation from the *wild* type structures (Fig 3B, 3D).

**Figure 2.**
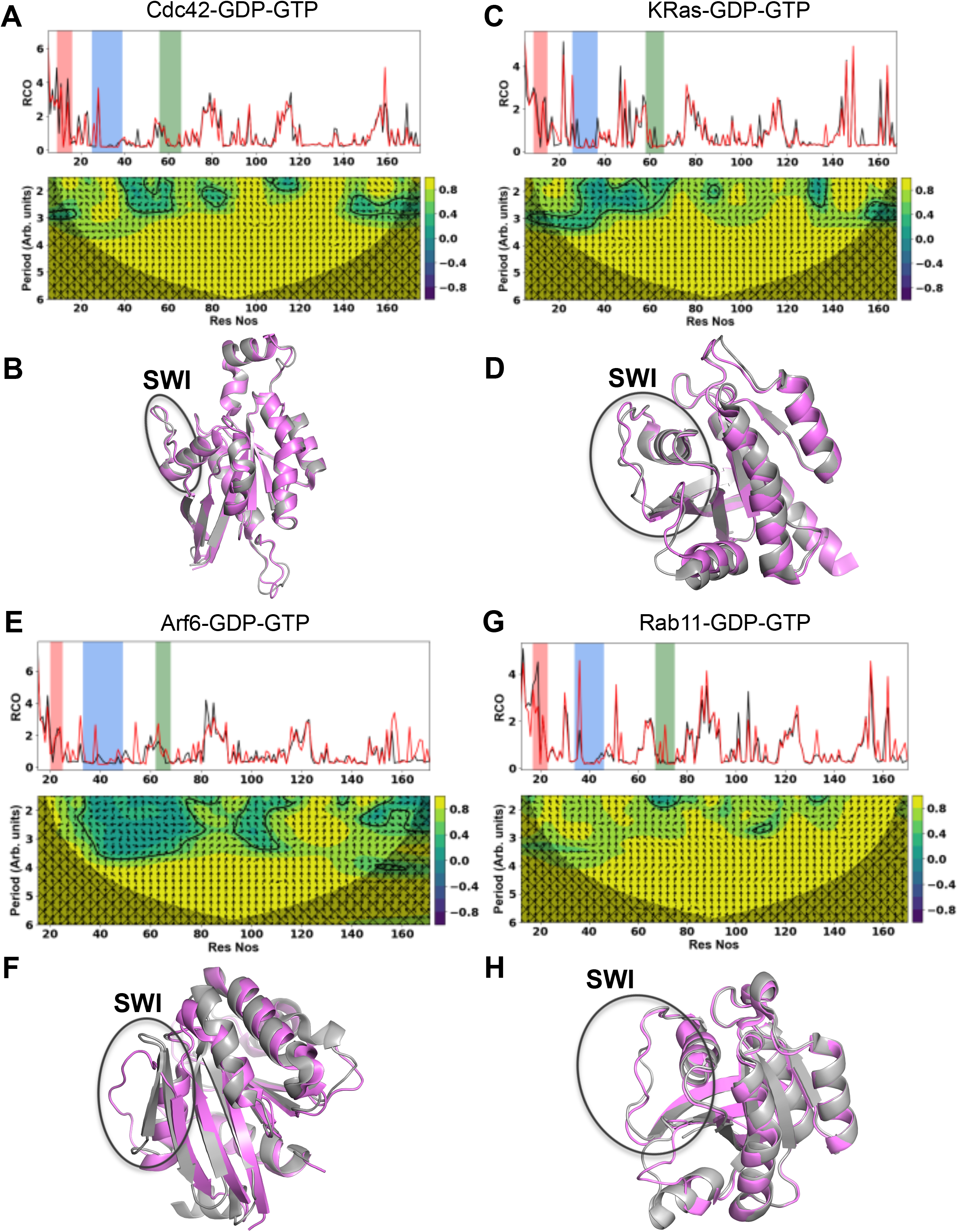
Residue Contact Order (RCO) and Wavelet coherence (WC) plots of GDP and GTP bound GTPase structures and their respective structure superimposition. Overlay of RCO plots and WC plots of *wild type* GTP and GDP bound structures of (A) Cdc42 (PDB CODES: 2QRZ & 1AN0) (C) KRas (PDB CODES: 6GOD & 4OBE) (E) Arf6 (PDB CODES: 2J5X & 1E0S) (G) Rab11 (PDB CODES: 1OIW & 1OIV). (B), (D), (F), (H) show the respective structure superimposition. In all the structure superimposition images GTP bound structures are shown in violet and the GDP bound structures are shown in grey and SWI region is highlighted with a black circle.

**Figure 3.**
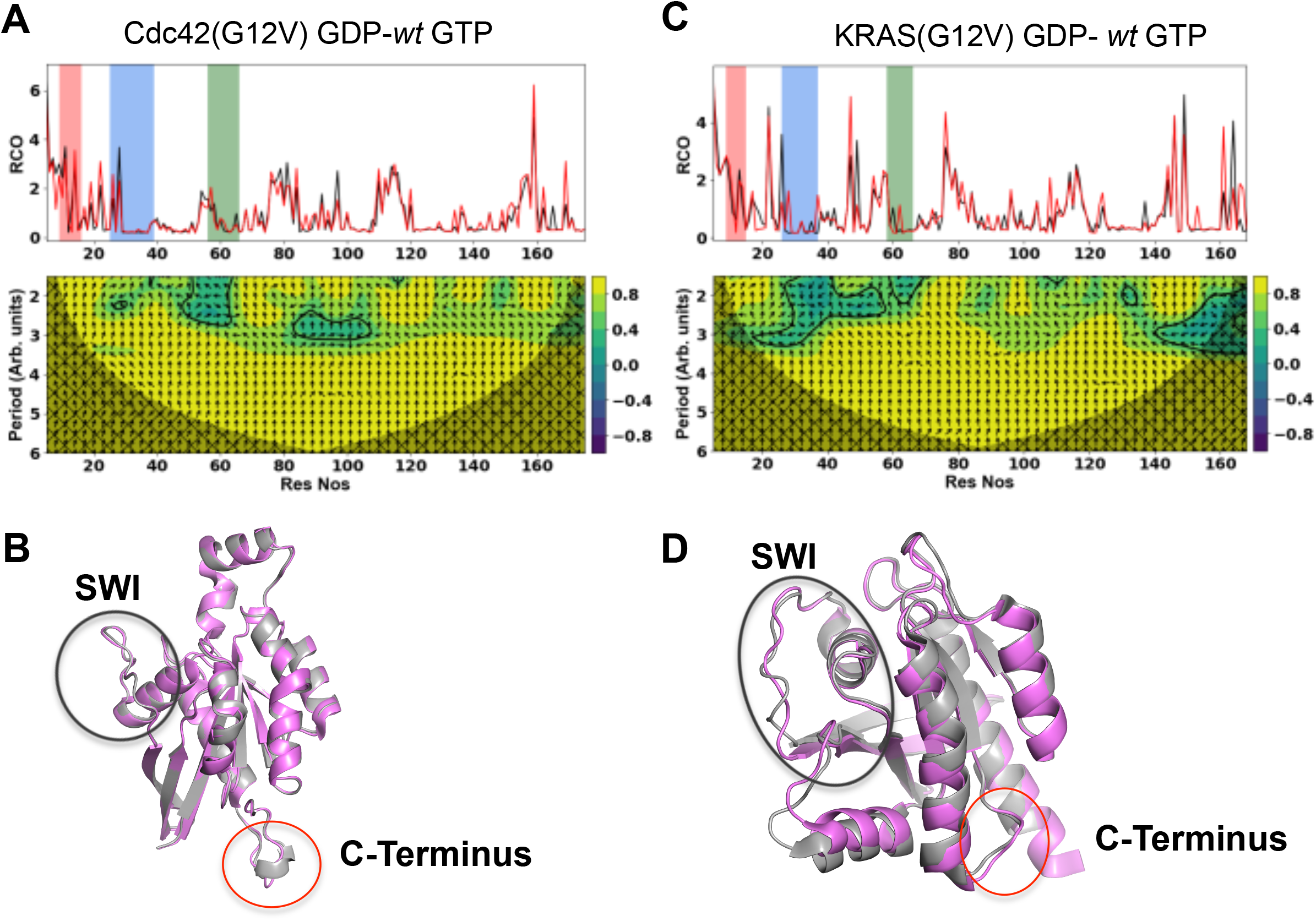
Comparison of RCO and WC of Cdc42 and KRas G12V GDP bound mutants with their GTP bound *wild type* structures. Overlay of RCO plots and WC plots of *wild type* GTP and G12V GDP bound mutant structures of (A) Cdc42 (PDB CODES: 2QRZ & 1A4R) (C) KRas (PDB CODES: 6GOD & 4TQ9) (B), (D) show the respective structure superimposition. The SWI region and C-terminus region showing significant conformational deviation are highlighted in black and red circles, respectively.

Several MD simulation based studies, involving principal component analysis (PCA), quantum mechanics/molecular mechanics (QM/MM) & network analysis suggested that there are no unique conformational signatures of SWI & SWII regions to define the functional states of the GTPases (Kumawat et al, 2017; Pálfy et al, 2019).

By and large, structural and MD simulation studies suggest that there are distinct ensembles of SWI & SWII conformers, which could be correlated with the functional states of the Ras superfamily GTPases and mutations in P-loop and switch regions affect the equilibrium of ensembles to alter the switching mechanism (Gorfe *et al*, 2008; Kapoor & Travesset, 2014; Li *et al*, 2018b; Muraoka *et al*, 2012; Prakash & Gorfe, 2013; Shurki & Warshel, 2004; Chandrashekar *et al*, 2011; Pálfy *et al*, 2019). Similar attempts have been made by analyzing the sequences of GTPases, employing Statistical Coupling Analysis (SCA), which has shown that evolutionarily conserved residues act at the functionally important regions map in a structurally connected network(Bandaru et al, 2017; Rivoire et al, 2016; Teşileanu et al, 2015). Altogether, MD and SCA studies suggest an allosteric model where the coupling of different structural regions dictates the dynamics and hence the functioning of the GTPases and aberrations in the coupling due to mutations would result in their abnormal functioning. However, these techniques do not clearly bring out the unified *modus operandi* of GTPases, manifested in terms of the structural allostery. In this paper, we have developed novel wavelet based tool (Fig 1B) to address questions like how the unified “loaded-spring” mechanism manifest across the Ras superfamily, despite exhibiting significant sequence and structural divergence? And how oncogenic mutations alter the loaded-spring mechanism?

Continuous wavelet transformation (CWT) has emerged as a powerful tool for feature extraction purposes, which can analyze localized and intermittent oscillations in the time series (Lee & Choi, 2019; Sato et al, 2018). Coherence analysis of wavelet transform of two signals, known as wavelet coherence analysis (WCA), simultaneously provides information on the relative changes between the signals and occurrence of these changes in the time domain with reasonable resolution. In the current study, we have applied WCA by mapping the 3D structures of proteins into 1D signals to unravel the co-related structural differences and gain functional insights.

Using this novel approach, we have analyzed the conformational variations in the Ras superfamily GTPases upon change in their bound nucleotide. Our results show that indeed there is a unified switching mechanism in the Ras superfamily, which manifests as coupling between wavelet coherence phase angles corresponding to the SWI & SWII regions. We tested the statistical significance of the observed correlation by populating the SWI and SWII conformers through MD simulation. These tests support our observations obtained from the comparison of just two (GTP and GDP) bound structures.

## Methods

### Conversion of 3D protein structures to 1D signal

Mathematically, a signal is nothing but changes in space or time, in some cases both. For example, an electrocardiogram (ECG) is a representation of a signal, which shows the electrical impulse passing through the heart as a function of time. Determination of fluctuations, such as discontinuities and breakdown points, in the signal will be of prime importance in delineating the information carried by the signal. Mathematical tools like Fourier or Wavelet transform (WT) aid in dissecting the meaningful information from the signals, which are otherwise not easily detectable. For instance WT has proved to be a powerful tool in accurate detection of QRS complex, T and P segments of ECG (Ivanov *et al*, 1999; Goldberger *et al*, 2002). These parameters determine how long the electrical wave takes to pass through the heart and pace of the wave to travel from one part of the heart to the other. This type of analysis provides information on the patho-physiology of the heart. Fourier Transform of any signal would give the frequency component present in it. Cardiac arrhythmias clearly reflect in the frequency based features of the ECG and have thus acted as a potential analytical tool for ECG. However, one of the major limitations of Fourier Transform is that the time resolution is lost in the frequency representation of the signal (Fig S2). In simpler words, from simple Fourier Transform, it is difficult to know at what time point in the signal did the frequency change. Short term Fourier Transform (STFT) provides some recourse to this problem but suffers from major drawbacks (Dremin & Nechitailo, 2007). The supplementary section elaborates further on this discussion.

The advantages of WT are that it may decompose a signal directly according to the frequency domains, which would be then represented in the time domain (Fig S3). Thus, both time and frequency information of the signal is retained. In simpler words, from WT one can know with reasonable resolution “when and how much” fluctuation occurred in the signal. Here, we have used WT techniques to determine both the quantum and the location of conformational changes in proteins, occurring due to change in their functional states. Proteins are the linear polymers of amino acids with distinct three-dimensional structures which define their function. Thus, three-dimensional structures of proteins can be looked as spatial changes of amino acids along the length of the ploy-peptide chain. For many class of proteins their functional states, like substrate/ligand bound-unbound & active-inactive states, could again be viewed as motions introduced in their polypeptide chains. To apply WT techniques on proteins we first transformed the three dimensional (3D) structure of the protein to a one dimensional signal (1D)(Chernick, 2001) by plotting the Residue Contact Order (RCO) for each of the residue against their respective positions. Mathematically, RCO of the *i*^th^ residue is defined as (Chen & Chaudhari, 2006; Kihara, 2005)

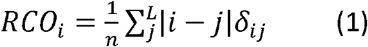

where n is the number of contacts between residue *i* and the others (*j*) belonging the polypeptide chain of length *L*, which lie within 3.5 Å of the *i*^th^ residue. Previous reports have defined RCO by excluding the contacts between the immediate neighbors of the residues (Chen & Chaudhari, 2006; Kihara, 2005). However, to generate a continuum model we have included the immediate neighboring residues, as well, for the calculations. RCO captures the spatial separation between the residues, which may not have covalent interactions, hence, encodes long range interactions present in the protein structure (Chen & Chaudhari, 2006; Kihara, 2005).

### Wavelet transformation of RCO signal of the proteins

The wavelet transform decomposes a signal as a combination of a set of basis functions, obtained by means of dilation (*s*) and translation (*τ*) of a proto type wave *φ*_0_ called as mother wavelet. As a characteristic, it has sliding windows that expand or compress to capture low- and high-frequency signals, respectively (Chernick, 2001). CWT of a signal *X(t)* is defined as

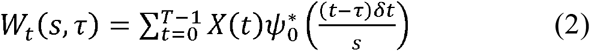

Where 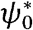 indicates complex conjugate of *ψ*_0_. s represents the scale, by varying which the wavelet can be dilated or contracted. By translating along localized time position one can calculate the wavelet coefficients *W*_*t*_(*s*, *τ*) which describe the contribution of the scales, to the time series *X(t)* at different time positions *t* (Cazelles *et al*, 2008)

The mother wavelet, which forms the orthonormal basis for the transformation, could be of different forms. For the current study, we have use Morlet mother wavelet, which is defined as

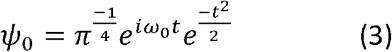

Wavelet coefficients are normalized to unit variance in order to allow direct comparisons of the wavelet coefficients. The local wavelet power, which describes how the special range of interaction of the residues in the structure varies along the residue positions, can be computed from wavelet coefficients as

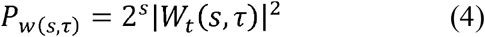

If the signal has finite length, as in the present case, wavelet transformation of it introduces zero padding effects. These are the artifacts present in the transformation that get more pronounced at the ends of the signal. Hence the wavelet power spectrum at the edges, in the region known as cone of influence (COI), is omitted for the analysis (Cazelles et al, 2008). The COI is demarcated in Fig 2A, 2C, 2E, 2G, 3A, 3C and S6.

### Wavelet coherence analysis

Wavelet coherence analysis provides insights on the variations between two signals. As mentioned earlier, wavelets are useful in delineating the “when and how much” of the variations between signals with reasonable resolution. The strength of the covariation between two signals (here conformational changes between two structures of a given protein in different states), represented as, *x(t)* and *y(t)* signals, can be quantified from the Wavelet coherence (WC) analysis. WC can be obtained from their cross wavelet transform (*w*_*xy*_), defined as

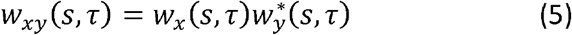

Where *w*_*x*_ is the CWT of the first structure [*x(t)*] and 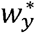 is the complex conjugate of CWT of the second structure [*y(t)*]. The power of the cross-spectrum is modulus of the CWT, |*w*_*xy*_(*s*, *τ*)|. The statistical significance of wavelet coherence is obtained by using Monte Carlo randomization techniques (Grinsted et al, 2004). The direction of the covariation between the signals can be obtained from the wavelet coherence phase (WCP), which can be calculated from the real, *R*(*w*_*xy*_(*s*, *τ*)), and the imaginary, *I*(*w*_*xy*_(*s*, *τ*)),parts of the cross wavelet transform as

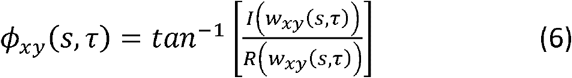

In the present situation, WCP provides propagation of the conformational variation for each of the residues as a function of their position in space. To quantify the net phase for a particular region a coherence weighted vector sum of all the phases, corresponding to the residues in that region, was obtained as

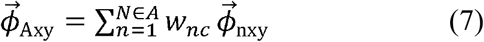

where *1….N* are the residues from the region *A* and 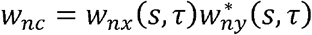, strength of the coherence at the n^th^ residue.

The algorithm is represented as a flowchart in Fig 1B and it is realized as a C program and a Python script. Both can be accessed on our GitHub page: https://github.com/drkakulkarni/WaveProtein. To perform DOCRR and MGC test we used the Hyppo software package (https://hyppo.neurodata.io/contributing.html).

## Results and Discussion

### Interpretation of Wavelet Transform of protein structures

Residue Contact Order (RCO) is a one-dimensional map of a three-dimensional structure that essentially represents change in the environment of a particular amino acid as a function of structural modulations. RCO captures both short-range and long-range interactions in a protein structure. Perhaps, this is one of the best one-dimensional structural representations of globular proteins. However, for extended protein structures such as collagen and large helical transmembrane proteins, RCO representation may appear to be skewered. The choice of distance cut-off in calculating the RCO plot is also critical in capturing the structural modulations. For example, 2.5Å cut-off would largely capture the short-range interactions, localized to the neighboring amino acids. On the other hand, a higher 10Å cut-off would largely mask the local modulations. In this study, we found 3.5Å cut-off for calculating RCO to be appropriate to map the three dimensional structure (Fig S4). This cut-off was found to work well with other globular proteins like Lysozyme and Heamoglobin (Fig S4B, S4C). Next, we tried to interpret the structural significance of the Wavelet Transform corresponding to RCO.

Prior to this, we obtained the Fourier Transform (FT) of the RCO as a first step to interpret wavelet transform. FT provides the frequency content of the signal. As evident from Fig S4, in the current context, frequency of the RCO signal corresponds to distinct classes of modulations present in the entire structure. For example, the 4^th^ residue may have two interactions with just the 10^th^ residue, which would result in RCO of the 4^th^ residue to be 3 (Eq 1). Similar “kind of” modulation could exist with other residues. For instance, if the 12^th^ residue has 4 interactions with only with 24^th^ then the RCO for this residue too would be 3. Example for other types of modulation could be the 32^nd^ residue interacting with the 36^th^ residue and having 10 contacts, the RCO for this residue would be 0.2. Thus, in this example, we see that two types of modulations (3 & 0.2) are exhibited by three different residues (at positions 4, 12, 32). The FT of the corresponding RCO would provide two peaks, corresponding to 0.2 and 3, however, the positional (residue) information, where the modulation is happening is lost. One more interesting observation from FT of RCO is that, the meaningful signal in the Fourier space exists only up to 10 frequency units (Fig S4 (green)). This is also true for other globular proteins such as, Hemoglobin and Lysozyme. Furthermore, FT provides hints on the optimal distance cutoff for calculating the RCO (Fig S4). The frequency information gets quenched for lower (2.5Å) distance as well as for very high (10Å) cut-off values (Fig S4).

As explained in the methodology section, Wavelet Transform captures both modulation (frequency) and location (time) information simultaneously with reasonable resolution. Therefore, wavelet transformation of RCO provides deconvoluted modulations at every residue in terms of scales reflecting different types of interactions (both long range and short range) (Fig S4). Thus, the co-relation between Wavelet Transform of two structures would provide relative change in the modulation at every amino acid and also the direction of propagation of the change. Here the quantum of change is represented by the magnitude of correlation (Eq 5) and the direction of change is represented by Wavelet Coherence Phase (Eq.no 6).

### Morlet Wavelet captures the backbone conformation of the protein

Details of the features that could be captured by the CWT depend on factors like the choice of the mother wavelet and scales chosen for the transformation (Fig S5 & S6). Here, we choose Morlet Wavelet (with ω_0_=6) and default scales (Fig S5) used in the python module of continuous wavelet transformation (https://pypi.org/project/pycwt/), to obtain the CWT of the protein 1D signal. Hence it is essential to validate the parameters chosen here are suitable for protein structural feature detection using CWT. To achieve this we calculated CWT of all known Rac and Cdc42 structures bound to GTP and GDP (Table S1) and sorted them using unsupervised *Kmeans* clustering (Supplementary methods). The outcome of this exercise would validate whether CWT used here are suitable enough to capture the conformational state of the GTPases. For this purpose, only main chain atoms of the proteins were used to calculate the RCOs and subsequent CWTs, as this would be sufficient to delineate the backbone conformation and at the same time mask the deviations in the structures due to mutations present in them. Our analysis show that the parameters used in calculating CWTs are optimal in capturing the conformational states of GTPases, as *Kmeans* clustering sorted the GTP and GDP bound structures with more than 80% efficiency (Table S2).

### Cross wavelet phase relationships show unique structural coupling between SWI and SWII regions of small GTPases

Next we asked whether cross correlation between the wavelet transformations (this is also known as wavelet coherence analysis), of GTP and GDP bound Ras GTPases provided tangible parameter to correlate the structure and functional states of GTPases. Since for Cdc42, KRas, Rab11 and Arf6 GTPases, complete (without breaks in the chains) GDP and GTP bound structures were available, they were chosen for the current analysis Fig 2 shows the wavelet coherence plots between the GTP and GDP bound *wild type* structures of these proteins.

Like the structure superpositions, even overlay of RCO plots by themselves do not show significant changes in the SWI, SWII and P-loop regions (Fig 2). Furthermore, even the autocorrelation analysis of RCOs, with a window of 5-10 residues, fails to conclusively show any correlation between the regions. However, RCO is a 1D projection of 3D structure, these plots would contain the information on the intrinsic dynamical alterations, which occur due to change in functional state of the protein. Hence to unravel these latent differential features of the structures, contained in the RCO, we performed Wavelet coherence analysis of the 1D signals of the protein structures (RCO plots). The advantage of using wavelet coherence analysis over the simple time domain coherence (also known as direct or time domain coherence), is that the former provides information on the quantum and location (residue position) of the relative changes, simultaneously, with reasonable resolution.

The y-axis of the WC plots corresponds to an arbitrary unit of period, which in the current context is a determinant of the range over which particular residue has ‘influence’ and the x-axis represents the residue numbers. In essence, the y-axis of the plot is a measure of 3D structural content of every residue mapped on 1D. The colour gradient of the plot is a measure of the strength of structural correlations between the GTP and GDP bound states, which is in the range of −1 (100% inversely correlated) to +1 (100% correlated). The arrows in the plot indicate the phase angles, measured in counter-clockwise direction. The arrows pointing right and left correspond to a zero angle (in phase) and 180°(out-of phase), respectively. Similarly, upward and downward pointing arrows correspond to part of the signal with 90° and 270° phase relationship, respectively.

Visibly, WC plots show changes in P-loop, SWI and SWII regions and these changes appear to be propagating from N to C terminal regions (Figs 2A, 2C, 2E and 2G). The quantum of change in these regions, as indicated by the WC-power (color of the region), appears to be same, which hints at the structural coupling between these regions. Another common feature in all of these plots is deviations at the C-terminus. Even though we have excluded about 5 residues at the N and C terminus of the structures, as they are floppy and disordered in many crystal structures, we see significantly correlated conformational differences at extreme regions. From these plots, absolute structural correlations of the individual regions in terms of WC power (colour gradient) appears to be challenging, as they are heterogeneous across the GTPases considered here. However, given the definition of the signal as 1D map of protein structure, the physical interpretation of arrows in the present context is propagation of relative conformational change.

Next, we asked whether WC analysis delineates the convergent switching mechanism, as there are no conserved conformational signatures between different GTPases families. To address this, we looked at the collective propagation of phase vectors corresponding to the P-loop, SWI and SWII regions to explore the signatures of conserved structural couplings that results in an orchestrated motion of these functional regions upon change in the bound nucleotide (Fig 4). For this, we calculated the resultant of the correlation weighted phase vectors belonging to these regions (Eqn 7), as the magnitude coherence provides the strength of change between the structures and phase vector provides direction of change. Since from Fourier space we can see that the meaningful structural modulations exists only upto 10 units of frequency (Fig S4), the phase vectors corresponding to the frequency levels greater than 10 were omitted for the calculations. Thus the resultant vector for a particular region (set of amino acids) provides a measure of net conformational change and the direction indicates the cumulative correlated change. From the definition of phase vectors, the net conformational propagation, in terms of angle between the vector corresponding to a particular region and the y-axis would be zero if there is absolutely no coordinated conformational change between the GDP and GTP bound states and a finite value of this angle serves as an index to compare the conformational changes across the GTPases.

**Figure 4.**
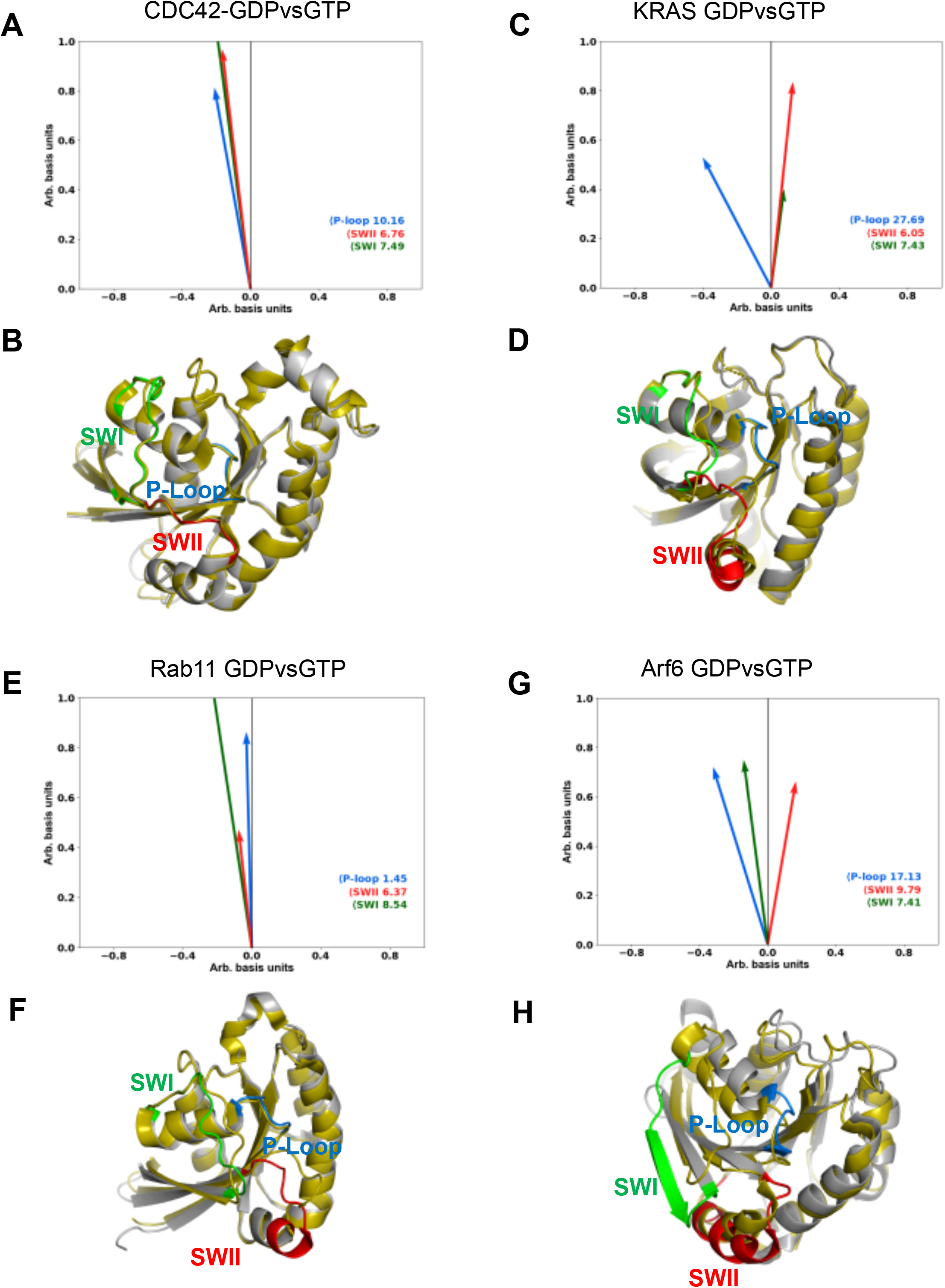
Wavelet coherence phase (WCP) vector plots of Ras superfamily members. **(A)** WCP Vector plot for *wild type* Cdc42 GTP& GDP bound structures and **(B)** their structure superimposition. **(C)** WCP Vector plot for *wild type* KRas GTP& GDP bound structures and **(D)** their structure superimposition. **(E)** WCP Vector plot for *wild type* Rab11 GTP& GDP bound structures and **(F)** their structure superimposition. **(G)** WCP Vector plot for *wild type* Arf6 GTP& GDP bound structures and **(H)** their structure superimposition. In the structure superimposition figures, GTP bound structures are shown in olive and the GDP bound structure is shown in grey. The PDB codes of the structures shown are as mentioned in Figs 2 and 3.

Interestingly, these vector plots (Fig 4) show that there is a well-defined phase relationship between SWI-SWII regions (Table 1, purple highlights), irrespective of the conformational changes seen in the crystal structures (Figs 4B, 4D, 4F, 4H). For almost all the structures considered here, the angle between Y-axis and the resultant phase vector corresponding to SWI and SWII regions lies in the range of 7 to 8 and 6 to 7 degrees, respectively. However the SWII phase vector of Arf6 (Table 1, grey highlight) exhibits deviation. Perhaps, this could be due to the role of additional regions in Arf switching mechanism, where the rearrangement of the switch regions is coupled to a variable N-terminal extension, which is auto inhibitory in the GDP-bound form and swings out to facilitate activation (Sztul *et al*, 2019). Further, to test whether the observed phase vector relationship is a consequence of the switching mechanism or it is an artifact of the methodology, we randomly morphed the SWI region in Cdc42^GDP^ structure and plotted the WC phase vector plot between the wild type GTP bound and the GDP bound morphed structures (Figs 5A and 5B). We clearly see that, in this instance, the conserved phase vector relationship is completely abolished, as the conformational changes introduced are random (Table 1, yellow highlight). Thus, the phase vector plots unambiguously bring out the conserved switching mechanism, which is not apparent in otherwise conformationally divergent Ras superfamily GTPases.

**Table 1.** Conformational deviations between the two different states of GTPases in terms of angles made by the resultant phase vectors corresponding to P-loop, SWI and SWII regions with y-axis. The GTPases that show similar trend in their SWI and SWII phase angles are highlighted in purple, the divergent Arf6 is shown in grey. The G12V mutants are highlighted in orange and the phase angles for the morphed Cdc42(GTP) is highlighted in yellow.

|  | P-Loop | SWI | SWII |
| --- | --- | --- | --- |
| *wtCdc42* (GTP)- *wtCdc42*(GDP) | 10.16 | 7.49 | 6.76 |
| *wtKRas* (GTP)- *wtKRas*(GDP) | 27.69 | 7.43 | 6.05 |
| *wtRab11* (GTP)- *wtRab11*(GDP) | 1.45 | 8.54 | 6.37 |
| *wtArf6*(GTP)- *wtArf6*(GDP) | 17.13 | 7.41 | 9.76 |
| *wtCdc42* (GTP)- G12V mutant *Cdc42*(GDP) | 13.34 | 14.23 | 6.26 |
| *wtKRas*(GTP)- G12V mutant *KRas*(GDP) | 12.70 | 8.96 | 10.39 |
| *morphed Cdc42* (GTP)- *wtCdc42*(GDP) | 13.14 | 21.29 | 11.19 |
| *wtCdc42*(**GDP**)- G12V mutant *Cdc42*(**GDP**) | 7.86 | 22.79 | 11.53 |
| *wtKRas*(**GDP**)- G12V mutant *KRas*(**GDP**) | 0.80 | 1.73 | 3.45 |

**Figure 5.**
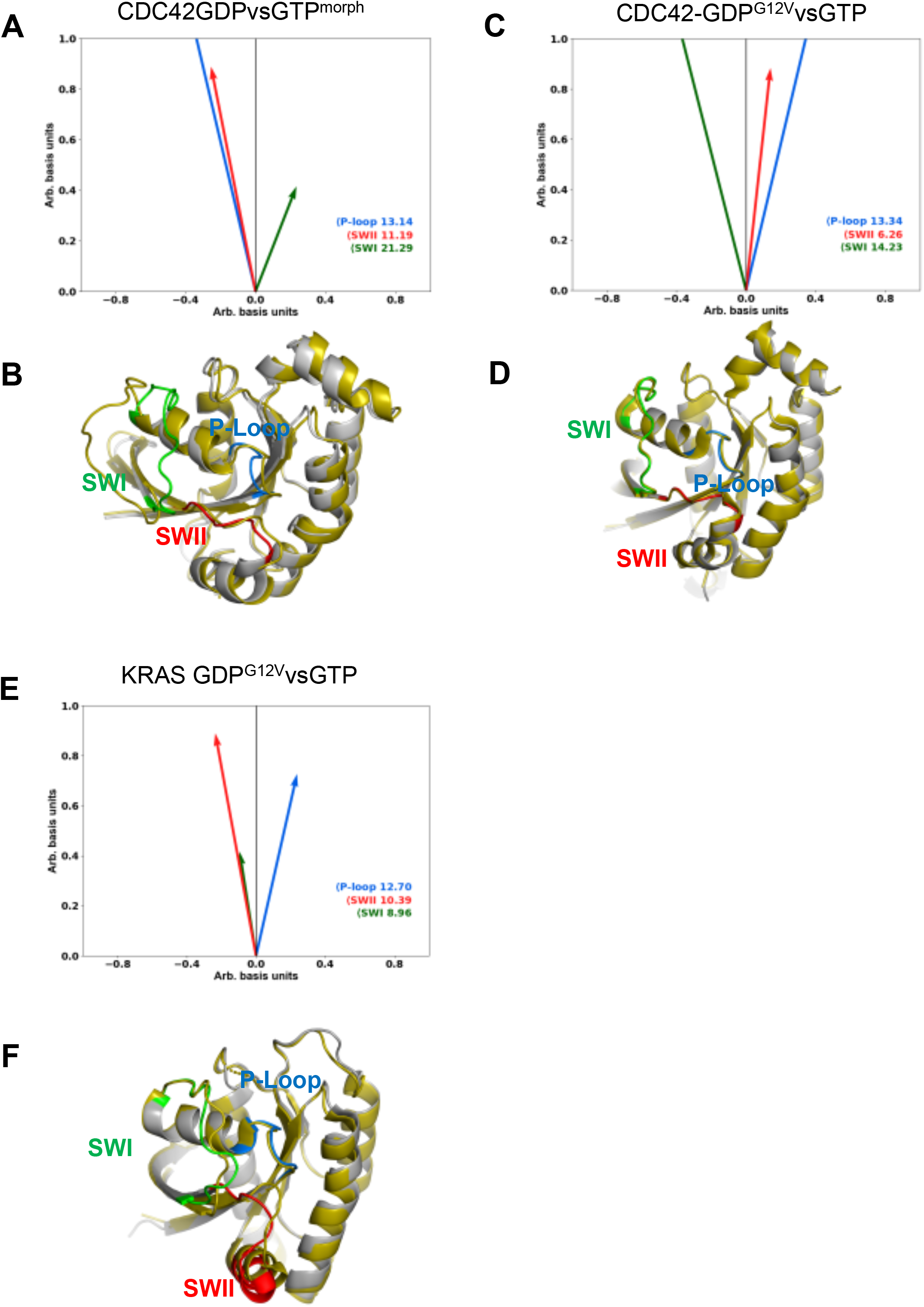
Wavelet coherence phase (WCP) vector plots of GTPases with aberrations. **(A)** WCP Vector plot for *wild type* Cdc42 morphed GTP& GDP bound structures and **(B)** their structure superimposition. **(C)** WCP Vector plot for *wild type* Cdc42 GTP & GDP bound G12V mutant structures and **(D)** their structure superimposition. **(E)** WCP Vector plot for *wild type* KRas GTP & GDP bound G12V mutant structures and **(F)** their structure superimposition. In the structure superimposition figures GTP bound structures are shown in olive and the GDP bound structure is shown in grey. The PDB codes of the structures shown are as mentioned in Figs 2 and 3.

### Oncogenic mutations in the P-loop disrupt the structural coupling between the functional regions of the GTPases

Although it is well known that oncogenic point mutations at 12^th^, 13^th^ (P-loop) and 61^st^ (SWII, residue numbering is with respect to the KRas structure) of the GTPases (Pantsar, 2020) abrogate their GTP hydrolyzing capacity, the exact role of the mutants in disrupting the GTP hydrolysis is not clear. Several studies, employing X-ray crystallography (Rudolph et al, 2008; Kawazu et al, 2013) and NMR (Smith *et al*, 2013; Pálfy *et al*, 2019; Chandrashekar *et al*, 2011) have shown that there is no significant structural differences between the *wild* type and mutant proteins, including their P-loop and switch regions (RMSD between the mutant and *wild* type structures is in the range 0.5 to 0.7Å). In the light of GTPase-GAP complex structures (Lu et al, 2016), using MD simulations and QM/MM calculations, it was proposed that disruption of GTP hydrolysis in the P-loop mutants (especially G12V/C mutants) could be due to increased flexibility at the nucleotide binding pocket, which would alter the GTP/GDP release rate (Khrenova *et al*, 2014; Sayyed-Ahmad *et al*, 2017; Pantsar, 2020). Thus, in the absences of major structural rearrangements (Figs 3B and 3D), just from structural analysis, it is difficult to explain how subtle changes at nucleotide binding pocket alter the intrinsic hydrolyzing capacity of the GTPases. The problem in deriving a generalized mechanism gets even more compounded due to structure and sequence divergences amongst the different family of GTPases at the nucleotide binding pocket. Since wavelet transformations are known to effectively separate individual signal components, we probed this problem with our WC tool. Comparison of WC phases corresponding to the *wild* type and G12V mutant of Cdc42 and KRas showed that the conserved phase coupling present in wild type proteins is disrupted in the mutant structure (Table 1, orange highlights, Fig 5). However the extent of change is varies between Cdc42 and KRas. As SWII region harbors the catalytically important Q61 residue and disentanglement in the P-loop-SWII coupling would essentially alter the Q61-solvent configuration, which could in turn affect the GTP hydrolysis. Thus WCA shows that the long-range interaction between functional regions of the GTPase are important for its biological function and minor aberrations in them would results in decoupling of the long range interactions leading to abrogation their function.

### Molecular dynamic (MD) simulations validate the WC phase coupling analysis

MD simulations have proved to be reliable mathematical tools to explore long-range interactions in proteins. Furthermore, MD simulations on many of the Ras family members have shown that the dynamics of switch regions is well correlated and can be deciphered form principal component analysis (Li *et al*, 2018a; Prakash & Gorfe, 2013; Kumawat *et al*, 2017). The current study involves comparison of just two structures, which may introduce bias in the results due to limited sampling of the conformational space. However small may be, a fraction of MD trajectories would contain conformer that resemble their native (initial) states. Therefore, we tested the correlation between the wavelet phase vectors, corresponding to the SWI and SWII regions, for different conformers generated through 100ns MD simulations. The MD simulations were performed on both *wild* type and mutant structures of KRas and Cdc42. The trajectories were sampled at every 200 ps to obtain 500 different conformers for which phase vectors were calculated using WCA tool (Fig S7). This exercise (Fig 6) clearly shows that even for a larger conformational space, the phase angles corresponding to SWI and SWII regions follow the trend observed with just two structures. To further enhance the confidence level of our findings, for one of structures (KRas GTP), we performed a very long 1μs simulation and sampled the trajectories at every 1000 ps. Interestingly, even for this larger dataset, the phase angles corresponding to SWI and SWII showed a similar correlation, as determined with only two conformers (Fig S8A). For Cdc42, we also performed reverse analysis, by using MD trajectories corresponding to the GDP bound and then comparing these with the GTP bound crystal structure. Even this exercise showed similar WC phase coupling between SWI and SWII regions. (Fig S8B). However, for mutants (Fig 6C, 6D), the joint distribution appears to be broadened. The PearsonR correlation values for the joint (bivariate) distribution of SWI and SWII WCA phases for the wild type proteins lie in the range of 0.4 to 0.8 (with p-values in the range 0.05 to 0.4) and for mutants it is 0.10 to 0.20 (with p-values in the range 0.1 to 0.9), suggesting modest linear correlation between the distributions, however, with poor statistical significance (Table 2). It is worth noting that these PearsonR values must be taken with caution, as the exact nature of the correlation (linear or non-linear) between the distributions is not clear. For non-linear data, distance correlation (DCORR) is widely used to bring out the intrinsic correlations (Lyons, 2013). Such “non-linear” covariance tests indicate the existence of correlations in the data in terms of the test p-values rather than the actual correlation values. Therefore, to check whether our data has any non-linear correlation, we performed the DCORR test on the phase angle distributions of SWI and SWII regions. Indeed, DCORR indicates that wild type phase angles of GTPases show significant non-linear-correlation, which gets disrupted in their mutants (Table 2). Although, DCORR has been shown to perform well in monotone relationships, but not so well in non-linear dependencies such as circles and parabolas (Shen *et al*, 2020). Recently, ‘Multiscale Graph Correlation’ (MGC) has emerged as a potential statistical test to unravel relationship and also the nature (geometry) of the relationship underlying multi-dimensional and/or nonlinear data (Shen *et al*, 2020; Vogelstein *et al*, 2019). Therefore we performed MGC test (Table 2, Fig S8) on our dataset to further validate the correlation between phase angles. For the wild-type protein the MGC p-value lies in the range of 0.004 to 0.009, with an exception for KRas short (100 ns) MD simulation data. On the contrary, mutant MD data indicate poor correlation between the SWI and SWII phase angle distribution with MGC p-values in the range of 0.8 to 0.7 (Table 2). These statistical tests suggest that, indeed the distributions of WCA phase angles have meaningful correlation for wild type proteins, which breaks down in the mutants that show aberrations in their switching action. This observation merits further investigations involving other GTPases. Thus, from these results it is evident that wavelet analysis has a potential to unravel intrinsic structural coupling by utilizing only two structures of the protein corresponding to the different functional states.

**Table 2.** P-Values from different correlation tests performed on the wavelet phase angles distributions corresponding to SWI and SWII regions. For Pearson, correlation values are shown in the parenthesis. PDB codes for the structures are same as given in Fig.1 and Fig. 2.

| Structure used in MD simulations | The counterpart crystal structure | Pearson (Correlation) | Distance Correlation | MGC |
| --- | --- | --- | --- | --- |
| *wtCdc42* (GTP) | *wtCdc42*(GDP) | 0.026(0.38) | 0.0001 | 0.009 |
| *wtCdc42*(GDP) | *wtCdc42* (GTP) | 0.432(0.44) | 0.0051 | 0.004 |
| *wtKRas* (GTP) | *wtKRas*(GDP) | 0.212(0.61) | 0.0028 | 0.020 |
| *wtKRas* (GTP) 1μs long simulation | *wtKRas*(GDP) | 0.05(0.78) | 0.0091 | 0.009 |
| G12V mutant *Cdc42*(GDP) | *wtCdc42* (GTP) | 0.13(0.11) | 0.2130 | 0.721 |
| G12V mutant *KRas* (GDP) | *wtKRas* (GTP) | 0.889(0.18) | 0.884 | 0.851 |

**Figure 6.**
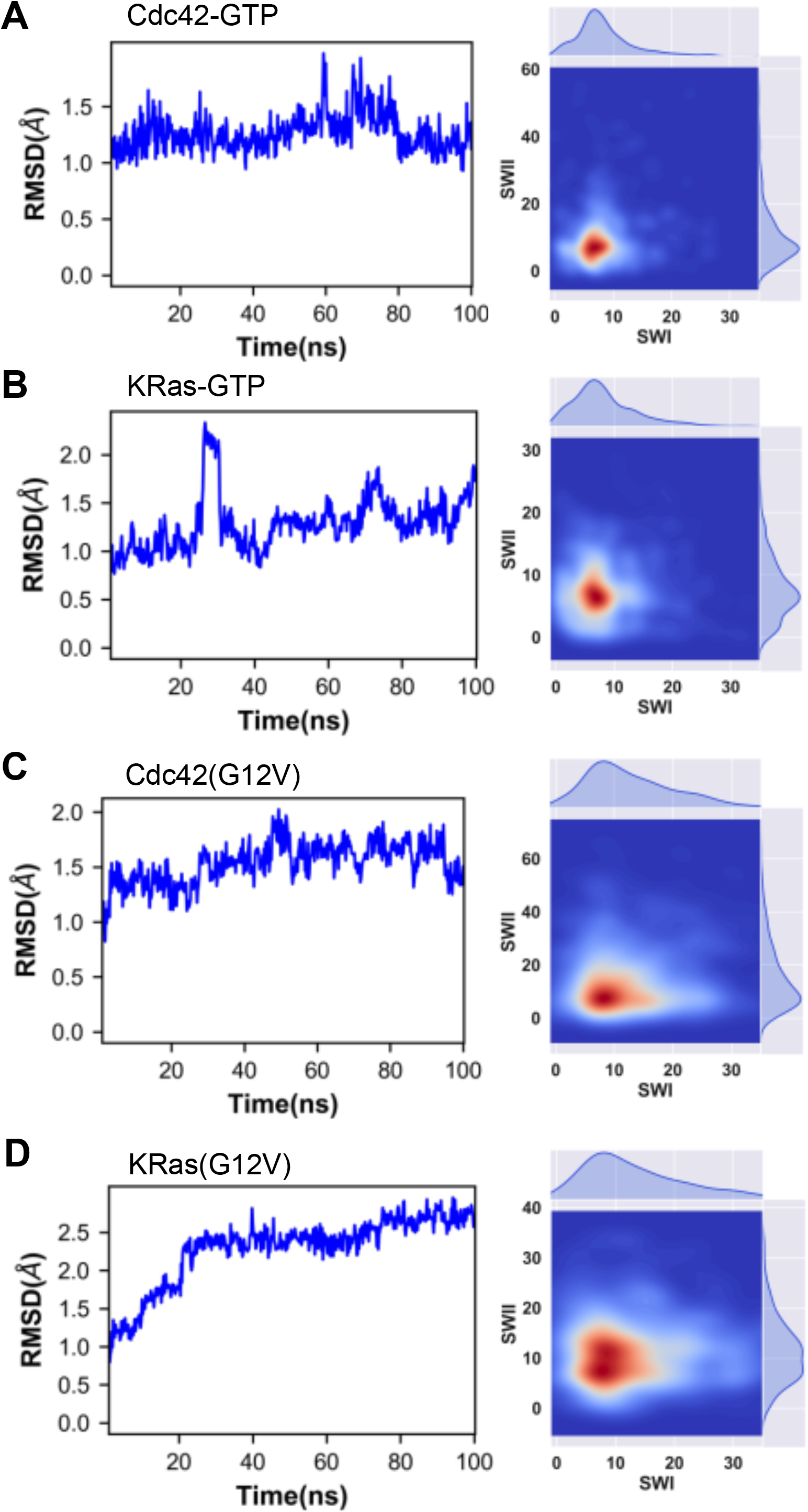
RMSD of MD trajectories of Cdc42 & KRas and bivariate distribution of SWI & SWII Wavelet coherence phase (WCP) angles. (Left) MD RMSD plots of GTP bound (A) Cdc42 (B) KRas and GDP bound G12V mutants of (C) Cdc42 and (D) KRas. The PDB codes of the structures shown are as mentioned in Figs 2 and 3. (Right) Bivariate distribution of SWI and SWII WC phase angles obtained from the MD simulation trajectories sampled at every 50ps with the static GTP or GDP bound forms of the respective GTPase.

## Conclusions

It is evident that function of proteins manifest in their structure and its dynamics, which is encoded in their sequence. However, subtle changes in the structure, either due to their functional states or mutations, cannot be easily comprehended from crystal structures. Perhaps, computational tools like MD simulations are found to be potential tools in probing protein dynamics. However, to derive generic inferences for a family of proteins it would require extensive simulation studies, which are computationally intractable and also suffer from convergence problem due to the non-linear nature of the method.

The wavelet-based methodology developed here is powerful enough to bring out the subtle dynamical aspects of proteins and it is computationally inexpensive. However, limitation of this approach is that it requires almost identical structures for comparison. Here by applying WCA, we have shown that Ras superfamily of small GTPases possess unique coupling in their functional regions despite exhibiting huge structural divergence in these regions. WCA provides a non-conventional interpretation of this coupling, which aids in unraveling the functional states of the protein and their aberrations due to mutations. Thus, this study could be a stepping-stone in delineating more complex dynamical aspects of protein structures and can be extended to other class of proteins, such as kinases, which exhibit conformational changes upon ligand binding.

## Acknowledgments

KK would like to acknowledge grant from Department of Biotechnology, India (BT/PR12502/BRB/10/1387/2015) and ZM would like to acknowledge fellowship from University Grants Commission, India. ASS would like to thank CSIR for fellowship (HCP0008)

## Author contributions

ZM and ASS performed data analysis. DS assisted with MD simulation analysis. KK designed the project, supervised overall work and wrote the paper with inputs from ZM, ASS and DS.

---

**Figure S1.**

**Active site of GDP and GTP bound KRas**.

**(A)** GDP bound KRas (PDB CODE: 4OBE) **(B)** GTP bound KRas (PDB CODE: 6GOD). The hydrogen bonding between γ-phosphate & Thr35 and γ-phosphate & Gly60 are shown with black dots and highlighted in a circle. GTP and GDP are indicated with yellow sticks and Mg2+ is indicated as a wheat colour sphere.

**Figure S2.**

**Advantages of Wavelet Transform over Fourier analysis of time signal.** (A), (P) A signal with the frequency component changing at different time points. In (A), frequency component increases with time while in (P), it is in the reverse order. (B) and (Q) represent the Fourier Transform of the signal in (A) and (P) respectively. (C) and (R) Wavelet Transform of (A) and (P) signals respectively. (D), (S) shows the effect of choice of scale on Wavelet Transform. Higher scales measure better frequency and truncation of lower frequency component is highlighted in a (D) red box, where Wavelet Transform on (A) is performed with higher scales.

**Figure S3.**

**Aspects of Wavelet Transform**. (A) The mother wavelet (Morlet in our study) is shown with different scales. Scaling elongates or contracts the mother wavelet. (B) (Left) Illustration of Wavelet Transform of signal shown in blue using the mother wavelet depicted in grey and black. The mother wavelet is translated along the time axis. (Right) The graphical representation of Wavelet Transform where each point represents the Wavelet Transform of the signal in terms of time and scale.

**Figure S4.**

**Relation between distance cut-off and frequency information corresponding to the RCO.** RCO (top) plots and their respective Fourier Transform (bottom) for (A) Cdc42 (PDB CODE: 1AN0) (B) Lysozyme (PDB CODE: 1L36) and (C) Hemoglobin (PDB CODE: 2DN2). The blue, green and red plots correspond to RCO calculated with distance cut-off of 2 Å, 3.5 Å and 10 Å respectively. As evident, the Fourier Transform of signals with 3.5 Å cut-off (green) show modulations upto 10 Hz, marked with a dotted line.

**Figure S5.**

**Wavelet Transform calculated on the RCO of Cdc42 (PDB CODE: 1AN0) with different scales and Morlet frequencies**. Encircled in the red box is the WT calculated with the parameters used for all analysis in this study. As seen, the chosen parameters bring out more features with less noise compared to others.

**Figure S6**

**Cone of Influence.** CWT of CDC42 GDP bound (PDB CODE: 1AN0). The COI is masked in black.

**Figure S7**

**Flowchart of WCP calculations.** Flowchart showing the algorithm for calculating the bivariate distribution of SWI & SWII Wavelet coherence phase (WCP) angles, employing MD simulation for GTP bound structures.

**Figure S8**

**RMSD of MD trajectories of Cdc42 & KRas, their bivariate distribution of SWI & SWII Wavelet coherence phase (WCP) angles and their MGC plots**.

(Left) MD RMSD plots of (A) GTP bound KRas (PDB CODE: 6GOD) (B) GDP bound Cdc42 (PDB CODE: 1AN0) (Center) Bivariate distribution of SWI and SWII WC phase angles obtained from the MD simulation trajectories sampled at every 200ps (for A) and 50ps (for B) with the static GDP or GTP bound forms of the respective GTPase. (Right) Their respective MGC test plots.

