## Supplementary material for "Wavelet coherence phases decode the universal switching mechanism of Ras GTPase superfamily": Supplimentary Methods

**Fundamentals of wavelet transform:**

Dissecting signals into smaller or sub-signals to extract features of interest is at the heart of signal processing. Mathematically, the objective is to decompose the signal and express it as a linear combination of other signals. These other signals are functions (eg. Sine, Cosine, Gaussian) that are often shifted and dilated.

i.e. if $X(t)$ is the signal and $\psi(t)$ is the other function then,

$$X(t)=\sum_{n} C_{n}\psi_{n}\left( t \right)$$

To put it simply, if a pianist simultaneously presses multiple keys of the piano to produce a particular sound, then, using signal processing, this signal can be decomposed in terms of sound produced by each of the keys pressed.

The numbers of functions depend on the type of signal and the functions used to decompose it. These functions are also known as bases functions and are generally orthonormal in nature. In Fourier Transform, these bases functions are sinusoidal in nature. Thus, Fourier Transform of any function provides information on the frequency content of the signal. However, the localization information of the frequency in the time domain gets diminished.

For example, in Fig S2, there are two waves (A, P) shown in the top panel. The first wave (Fig S2A) has lower frequency content in the beginning 2 seconds and higher frequency content later, on the contrary, the second wave (Fig S2P) is the opposite of first wave. However, the Fourier Transform of both waves (Fig S2 B, Q) is identical, as this method provides the frequency content present in the wave notwithstanding the occurrence of these frequencies in the time domain.

To obtain the localized frequency content, a Fourier Transform combined with a window $w\left( t-\tau\right)$ that moves over the signal is necessary. This type of Fourier Transform is termed as Short Time Fourier Transform (STFT). The primary limitation of STFT is that the resolution is bounded with the chosen window size. In simpler words, a smaller window would give better time resolution but compromise the frequency resolution and vice-versa. This problem is circumvented to a great extent in Wavelet Transform (WT).

In WT, the window is combined with the analyzing function and thus the information to be obtained depends on the analyzing function. This function is also known as the mother wavelet and it is not only translated along the signal but can also be dilated. The dilation of the mother wavelet (analyzing function) is known as scaling (s). The mother wavelets can be of different functions and in our current study we have used Morlet wave (equation 3 of main text) as the mother wavelet with scale 1. Figure S3A, shows Morlet function with two different scales. Morlet wavelet is found to be useful in determining the local changes (frequencies) and therefore used for the current study. From equation 2 of the main text, it is clear that Continuous Wavelet Transform (CWT) provides time and frequency information by convoluting the wavelet with the signal at different locations.

**Supplementary Methods**

**Unsupervised clustering of GTPase structures**

Python implementation of KReas image clustering was employed to cluster the wavelet transforms of the GTPase structures (Nelli & Nelli, 2018). The COI of Wavelet transformation spectrums of the structures were masked to avoid influence of the artifacts in sorting.

**MD simulation methods**

Molecular dynamics (MD) simulations were carried out using Desmond v3.6 (Shivakumar *et al*, 2010; Guo *et al*, 2010; Bowers *et al*, 2006), implemented in Schrodinger-maestro 2020 (Schrödinger Release, 2019). Simulation system was prepared by placing the GTPase in an orthogonal box with 10.0Å buffered distance in all the directions, after assigning bonds. The hydrogen atoms were added to the protein keeping pH 7.0 of the model. The system was solvated with TIP3P (Jorgensen *et al*, 1983) solvent atoms and neutralized by the addition of Na+/Cl- ions. Subsequently, the system was subjected to energy minimization followed by 2000 steps of conjugate gradient algorithm with 120.0 kcal/mol/Å convergence threshold energy under NPT ensemble. Finally, simulation of the system was performed for 100 ns. Trajectory recorded in each 200 ps. For KRas GTP, a longer simulation of 1 μs was performed and the trajectories were sampled at every 1ns. All the simulations, except the 1 μs KRas simulation, were performed in triplicates.

**Table S1.**

List of PDBs used

| Sl. No | Name of the GTPase | PDB Code | Remark | Organism | Reference |
| --- | --- | --- | --- | --- | --- |
| 1 | CDC 42 (F28L mutant) | 2ASE | Nucleotide free | Homo sapiens | (Adams & Oswald, 2006) |
| 2 | RHOA (Complex with RhoGEF) | 4D0N | GDP bound | Homo sapiense | (Azeez et al, 2014) |
| 3 | RAC1(complex with Plexin-B1) | 3SU8 | GTP bound | Homo sapiens | (Bell et al, 2011) |
| 4 | CDC 42 (complex with Bacterial Toxin Sope) | 1GZS | GTP bound | Homo sapiens | (Buchwald et al, 2002) |
| 5 | RAC2 (G12V mutant) | 2W2T | GDP bound | Homo sapiens | (Bunney et al, 2009) |
| 6 | RAC1 (complex with P-Rex1) | 5FI0 | Nucleotide free | Homo sapiense | (Cash et al, 2016) |
| 7 | CDC42 (complex with P-Rex1) | 5FI1 | Nucleotide free | Homo sapiense | (Cash et al, 2016) |
| 8 | RHOA (complex with GEF) | 3KZ1 | GTP bound | Homo sapiens | (Chen et al, 2010) |
| 9 | RHOA (complex with RhoGEF) | 6BCA | GTP analouge bound | Homo sapiense | (Chen et al, 2018) |
| 10 | RHOA (complex with RhoGEF) | 6BCB | GTP analouge bound | Homo sapiens | (Chen et al, 2018) |
| 11 | RAC1(complex with Trio) | 2NZ8 | Nucleotide free | Homo sapiens | (Chhatriwala et al, 2007) |
| 12 | KRAS | 6GOD | GTP bound | Homo sapiens | (Cruz-Migoni et al, 2019) |
| 13 | RHOA (complex with RhoGEF) | 6BC0 | GTP analouge bound | Homo sapiense | (Dada et al, 2018) |
| 14 | RAC1 (complex with RhoGEF) | 6BC1 | GTP analouge bound | Homo sapiense | (Dada et al, 2018) |
| 15 | RAC1 F28L mutant | 4GZM | GTP bound | Homo sapiens | (Davis et al, 2013) |
| 16 | RHOA (complex with the DH/PH fragment of PDZRhoGEF) | 1XCG | Nucleotide free | Homo sapiens | (Derewenda et al, 2004) |
| 17 | RAC1 | 5N6O | GDP bound | Homo sapiense | (Ferrandez et al, 2017) |
| 18 | RHOA (complex with RHOGAP) | 1OW3 | GDP bound | Homo sapiens | (Graham et al, 2002) |
| 19 | RAC1 (complex with RHOGDI) | 1HH4 | GDP bound | Homo sapiens | (Grizot et al, 2001) |
| 20 | RAC1 (T17N mutant) | 3B13 | Nucleotide free | Homo sapiens | (Hanawa-suetsugu et al, 2012) |
| 21 | CDC42 (T17N mutant) | 3VHL | Nucleotide free | Homo sapiense | (Harada et al, 2012) |
| 22 | RAC1 | 1MH1 | GTP analogue bound | Homo sapiens | (Hirshberg et al, 1997) |
| 23 | CDC 42 (Complex with RHOGDI) | 1DOA | GTP bound | Homo sapiens | (Hoffman et al, 2000) |
| 24 | KRAS | 4OBE | GDP bound | Homo sapiense | (Hunter et al, 2014) |
| 25 | KRAS G12V mutant | 4TQ9 | GDP bound | Homo sapiens | (Hunter et al, 2015) |
| 26 | RHOA | 1A2B | GTP Analogue bound | Homo sapiens | (Ihara et al, 1998) |
| 27 | RHOA | 3TVD | GTP bound | Rattus norvegicus | (Jobichen et al, 2012) |
| 28 | RAC1 (P29S mutant) | 3SBD | GTP bound | Homo Sapiens | (Krauthammer et al, 2012) |
| 29 | RAC1 | 3TH5 | GTP bound | Homo sapiens | (Krauthammer et al, 2012) |
| 30 | RHOA (complex with RhoGDI) | 5FR2 | GDP bound | Homo sapiense | (Kuhlmann et al, 2016a) |
| 31 | RHOA (complex with RHOGDI) | 5FR1 | GDP bound | Homo sapiense | (Kuhlmann et al, 2016b) |
| 32 | CDC 42(complex with DOCK7) | 6AJ4 | Nucleotide free | Homo sapiense | (Kukimoto-Niino et al, 2019) |
| 33 | RAC1 (complex with DOCK2) | 2YIN | Nucleotide free | Homo sapiens | (Kulkarni et al, 2011) |
| 34 | CDC 42(complex with GAP) | 5CJP | GTP bound | Homo sapiens | (LeCour et al, 2016) |
| 35 | Rab 28 | 3E5H | GTP bound | Homo sapiens | (Lee et al, 2008) |
| 36 | RHOA (Complex with RHOGDI) | 1CC0 | GDP bound | Homo sapiens | (Longenecker et al, 1999) |
| 37 | RHOA (Q63L mutant) | 1KMQ | GTP analogue bound | Homo sapiens | (Longenecker et al, 2003) |
| 38 | RAC1 (complex with P-Rex1) | 4YON | Nucleotide free | Homo sapiense | (Lucato et al, 2015) |
| 39 | ARF6 | 1E0S | GDP bound | Homo sapiens | (Ménétrey et al, 2000) |
| 40 | CDC 42 (complex with Cdc42GAP) | 1GRN | GDP bound | Homo sapiens | (Nassar et al, 1998) |
| 41 | Arf6 | 2J5X | GTP bound | Homo sapiens | (Pasqualato et al, 2001) |
| 42 | CDC42 | 2QRZ | GTP bound | Homo sapiens | (Phillips et al, 2008) |
| 43 | RAC1 (complex with Protein kinase ypkA) | 2H7V | GDP bound | Homo sapiens | (Prehna et al, 2006) |
| 44 | RAC1(complex with GEF-Vav1) | 2VRW | Nucleotide free | Homo sapiens | (Rapley et al, 2008) |
| 45 | CDC 42(complex with GEF Dbs) | 1KZ7 | Nucleotide free | Homo sapiens | (Rossman et al, 2002) |
| 46 | CDC 42 (Complex with GEF dbs) | 1KZG | Nucleotide free | Homo sapiens | (Rossman et al, 2002) |
| 47 | CDC 42 (G12V mutant) | 1A4R | GDP bound | Homo sapiens | (Rudolph et al, 2008) |
| 48 | RAC2 (Complex with RHOGDI) | 1DS6 | GDP bound | Homo sapiens | (Scheffzek et al, 2000) |
| 49 | RHOA | 1DPF | GDP bound | Homo sapiens | (Shimizu et al, 2000) |
| 50 | CDC 42(complex with Dbl exchange factors) | 1KI1 | Nucleotide free | Homo sapiens | (Snyder et al, 2002) |
| 51 | RAC1 (complex with Salmonella effector SptP) | 1G4U | GDP bound | Homo sapiens | (Stebbins & Galán, 2000) |
| 52 | RHOA (complex with RhoGDI) | 4F38 | GTP bound | Mus musculus | (Tnimov et al, 2012) |
| 53 | RAC1 | 6AGP | GDP bound | Homo sapiens | (Toyama et al, 2019) |
| 54 | RHOA | 1FTN | GDP bound | Homo sapiens | (Wei et al, 1997) |
| 55 | RAC1 | 1FOE | Nucleotide free | Homo sapiens | (Worthylake et al, 2000) |
| 56 | RAC1(complex with Pseudomonas Aeruginosa Exos Toxin) | 1HE1 | GDP bound | Homo sapiens | (Würtele et al, 2001) |
| 57 | CDC 42 (complex with GEF) | 2DFK | Nucleotide free | Homo sapiens | (Xiang et al, 2006) |
| 58 | CDC 42 (complex with Dock9) | 2WM9 | Nucleotide free | Homo sapiens | (Yang et al, 2009) |
| 59 | CDC 42 (complex with Dock9) | 2WMN | GDP bound | Homo sapiens | (Yang et al, 2009) |
| 60 | CDC 42 (complex with Dock9) | 2WMO | GTP bound | Homo sapiens | (Yang et al, 2009 |
| 61 | RHOA (complex with RhoGAP) | 5HPY | GDP bound | Homo sapiense | (Yi et al, 2016) |
| 62 | RAC3 | 2G0N | GDP bound | Homo sapiens | To be Published |
| 63 | Rab 28A | 2HXS | GDP-3'P-Bound | Homo sapiens | To be Published |
| 64 | RAC3 | 2IC5 | GTP analogue bound | Homo sapiens | To be Published |
| 65 | CDC 42 (T35A mutant) | 2KB0 | Nucleotide free | Homo sapiens | To be Published |
| 66 | RHOA (complex with RhoGAP) | 3MSX | GDP bound | Homo sapiens | To be Published |
| 67 | CDC42 (complex with RACGAP) | 5C2J | GDP bound | Mus musculus | To be Published |
| 68 | RHOA(complex with RACGAP) | 5C2K | GDP bound | Homo sapiens | To be Published |
| 69 | RHOA (complex with ARHGEF) | 5JHG | Nucleotide free | Homo sapiense | To be Published |
| 70 | RHOA (complex with ARHGEF) | 5JHH | Inhibitor bound | Homo sapiens | To be Published |
| 71 | CDC 42 | 1AN0 | GDP bound | Homo sapiens | To be published |

**Table S2.**

Sorting of GTP and GDP bound structures by Kmeans clustering

|  | **Prediction Efficiency (%)** | **False Positive (%)** |
| --- | --- | --- |
| **GTP** | 85.7% | 28.5% |
| **GDP** | 83.3% | 8.93% |
