## Supplementary figures and images for "Wavelet coherence phases decode the universal switching mechanism of Ras GTPase superfamily"

### Supplimentary Figures

#### Figure S1

A

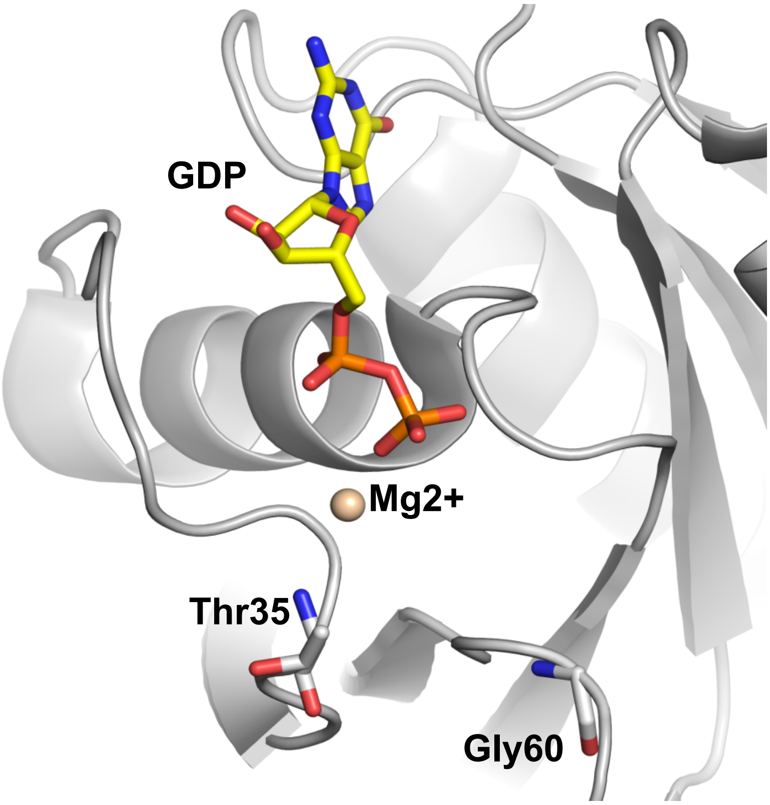


B

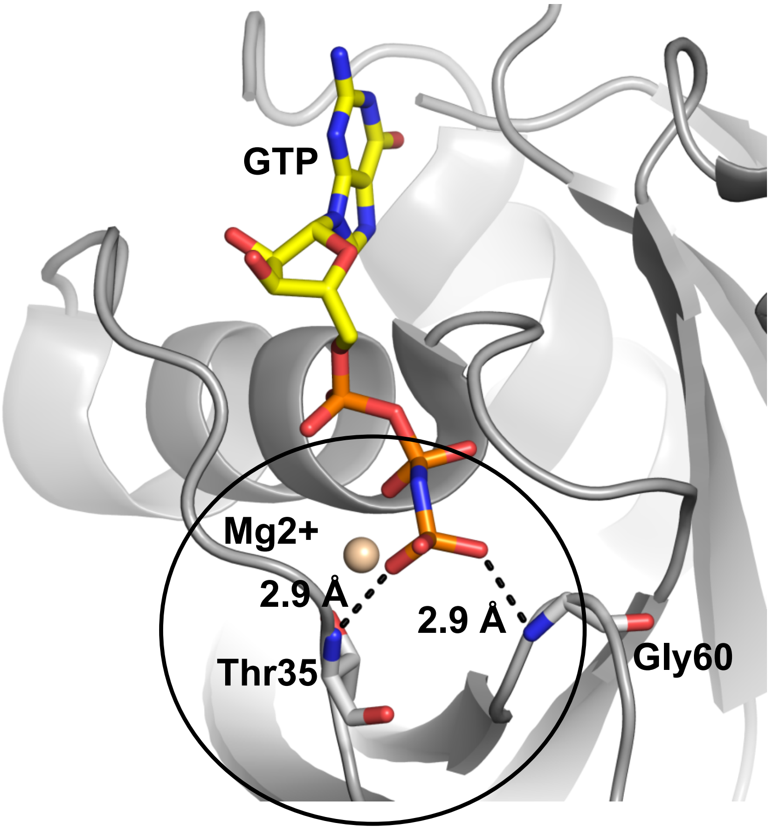

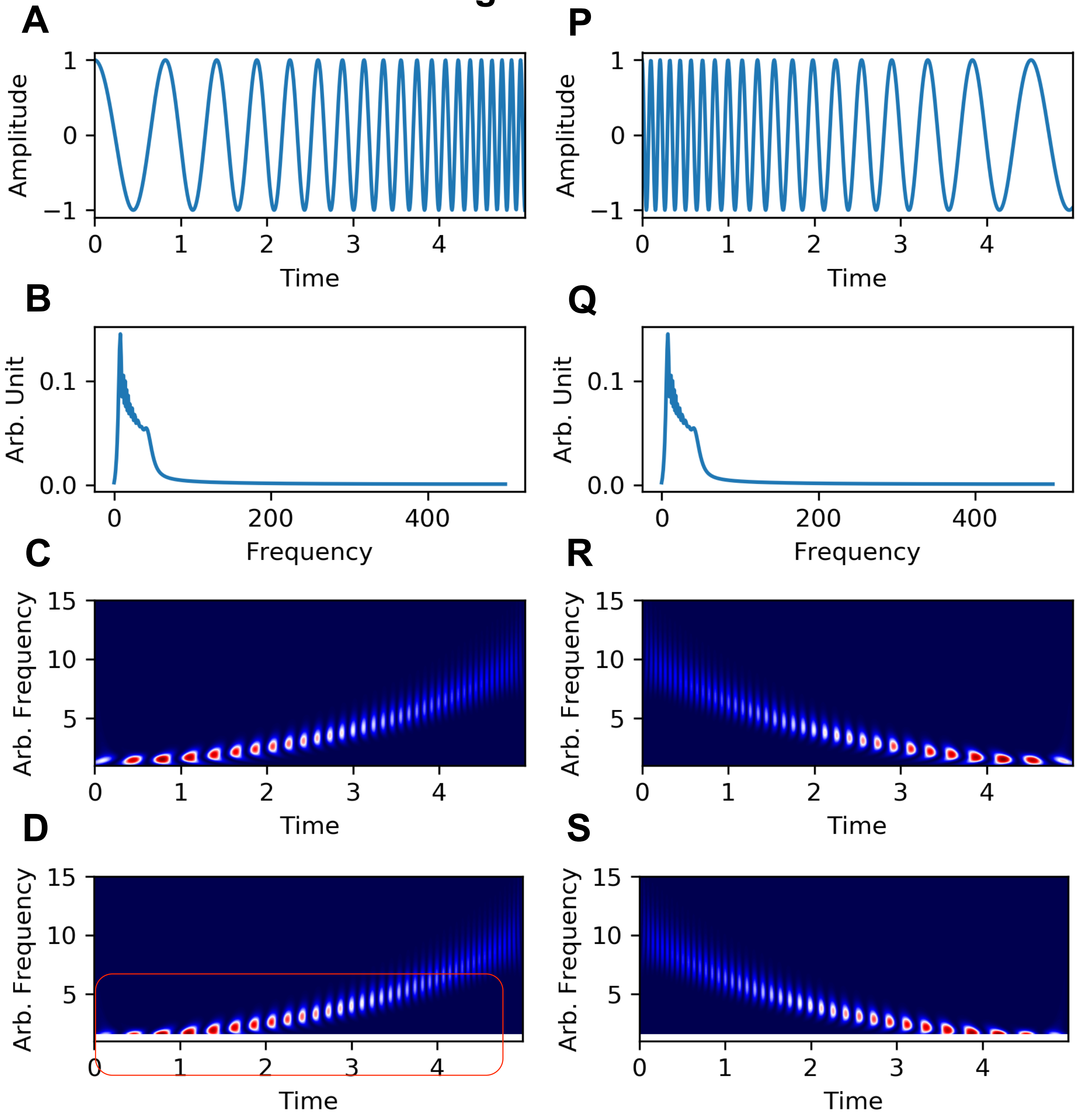

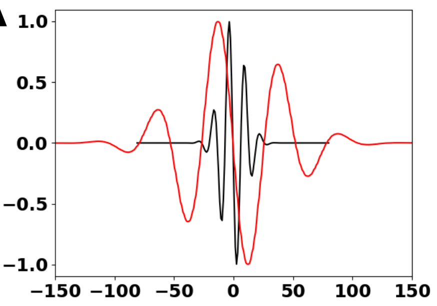

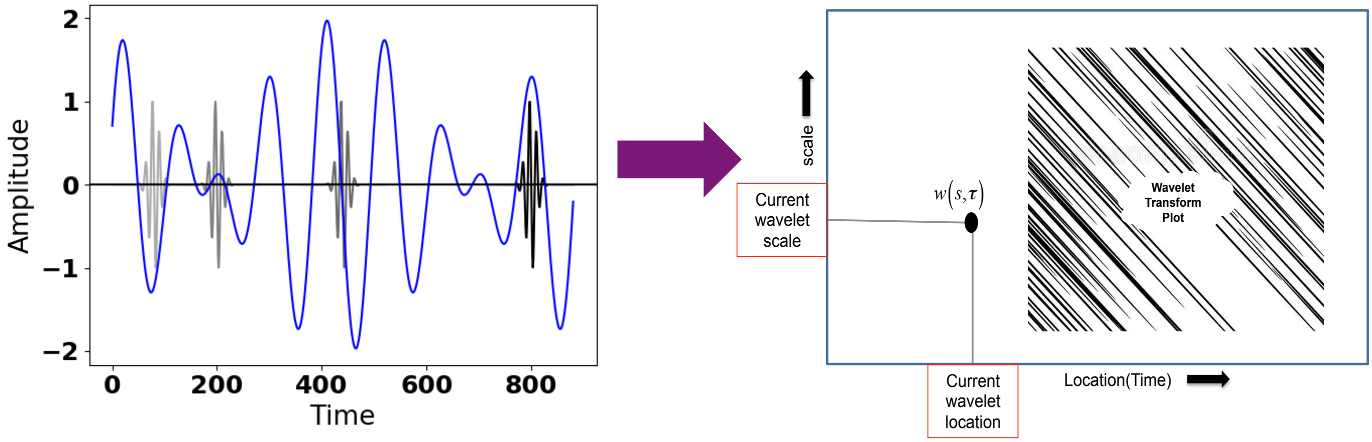

#### Figure S4

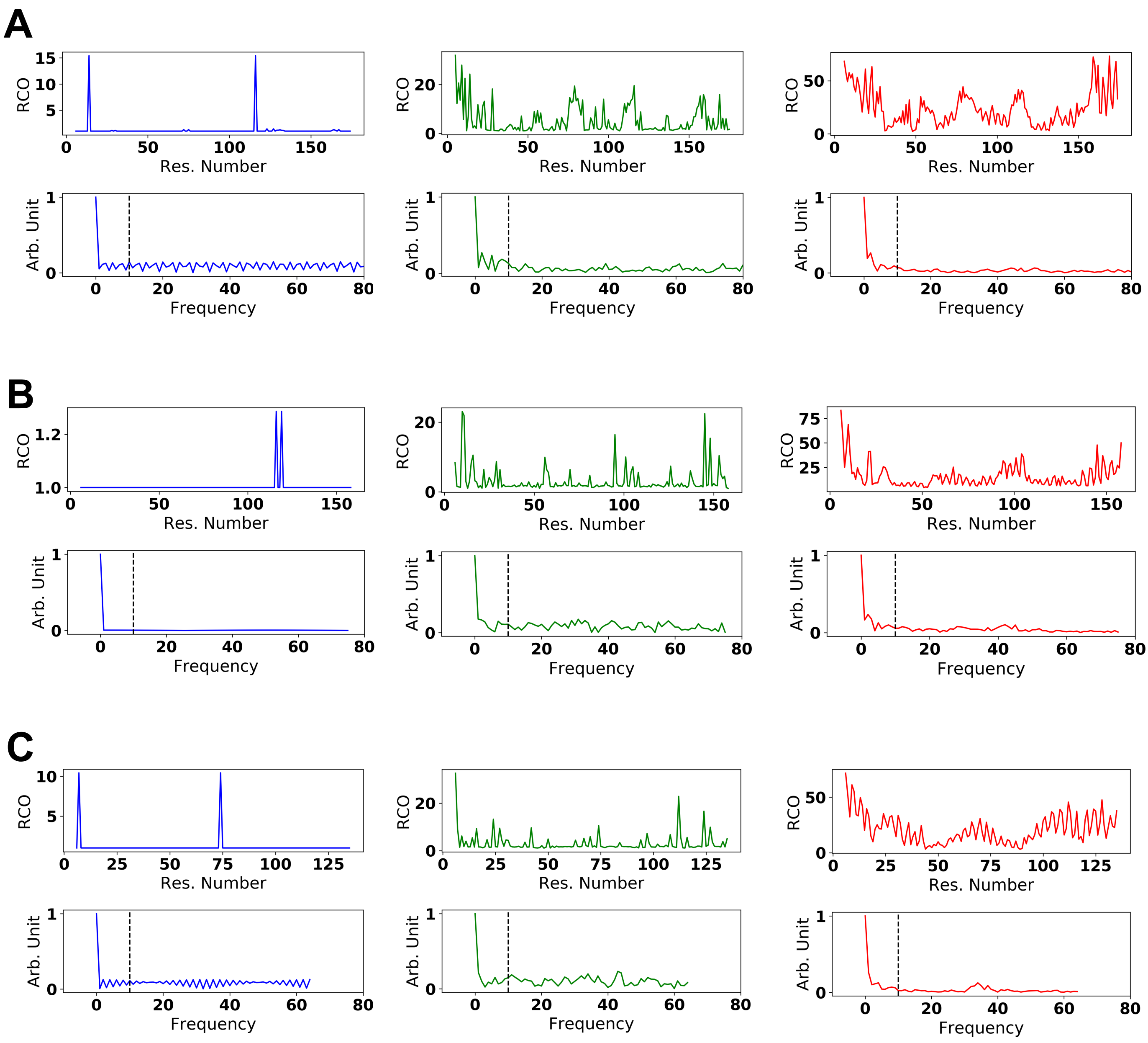

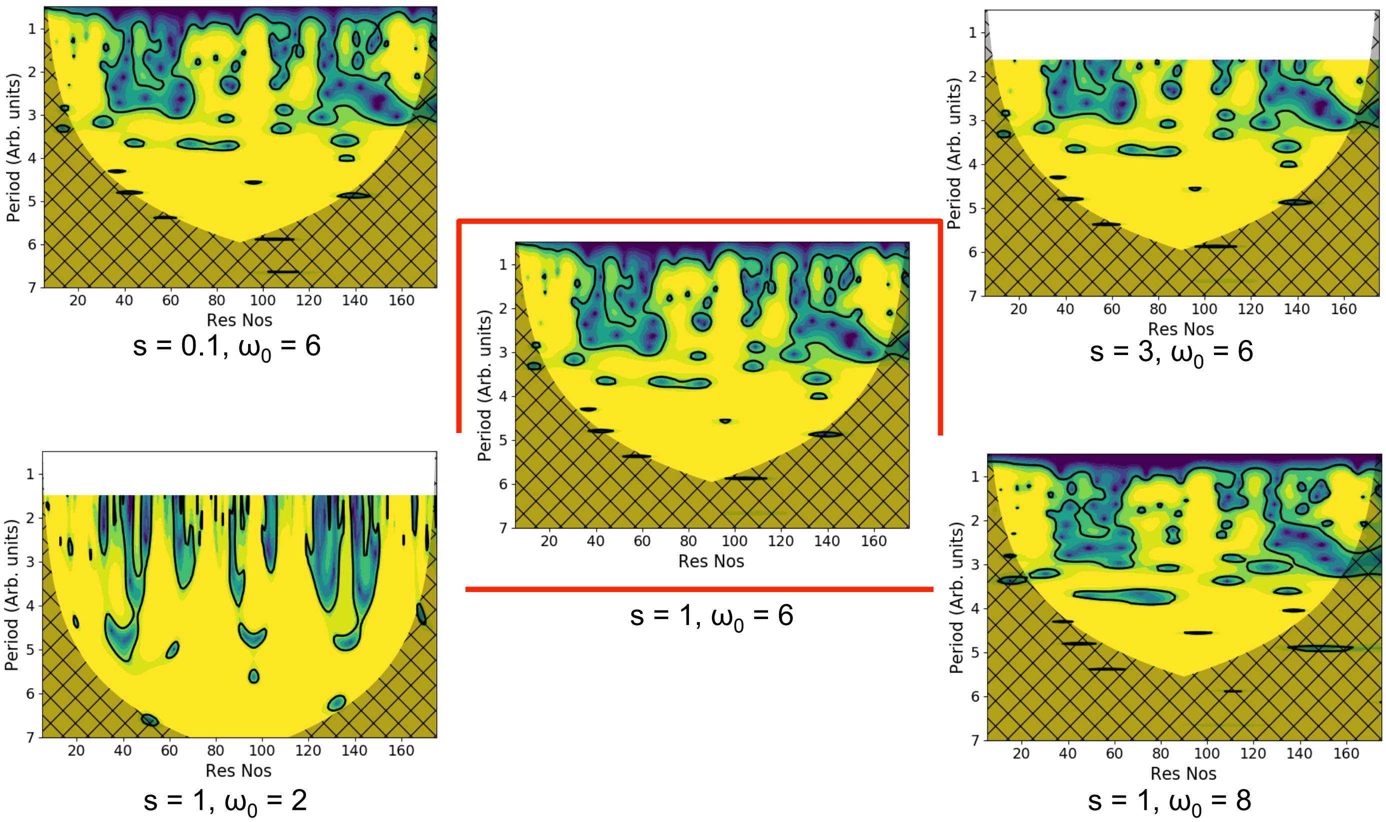

#### Figure S6

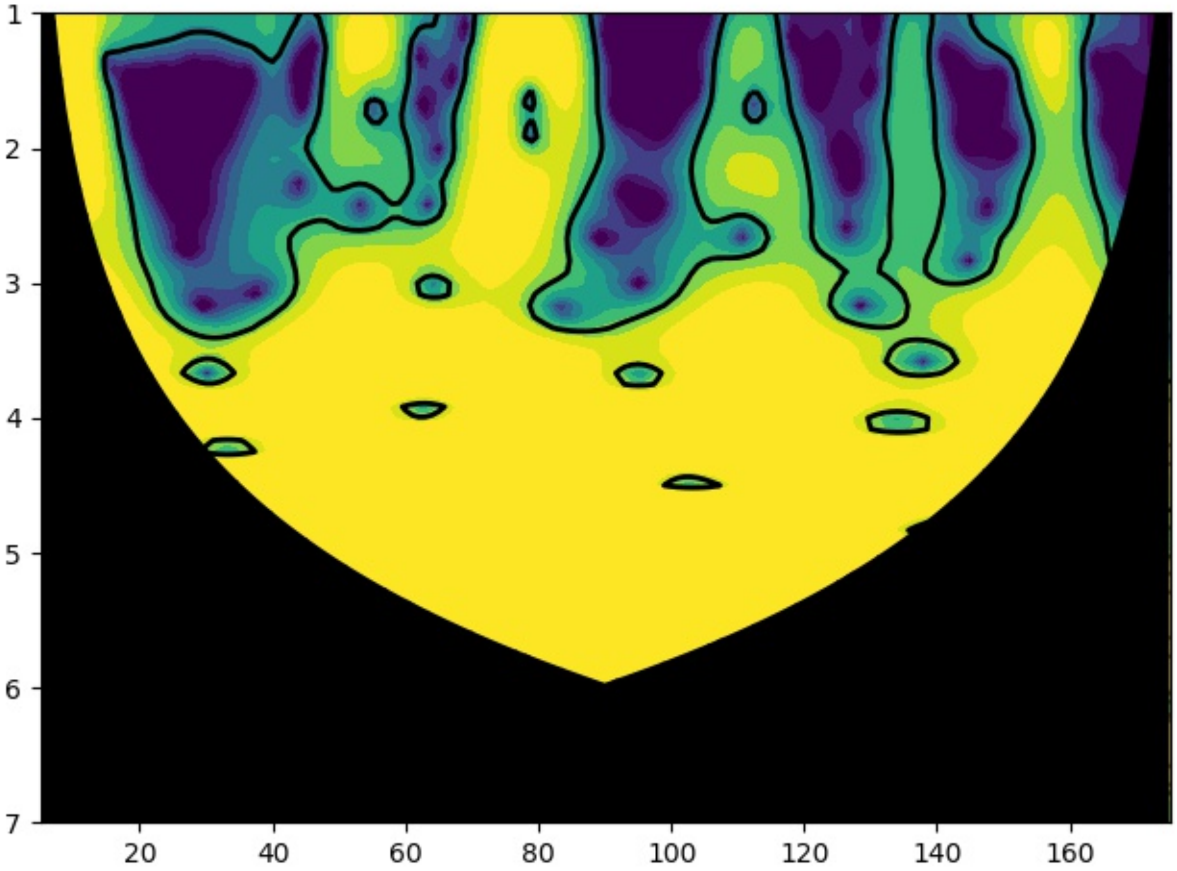

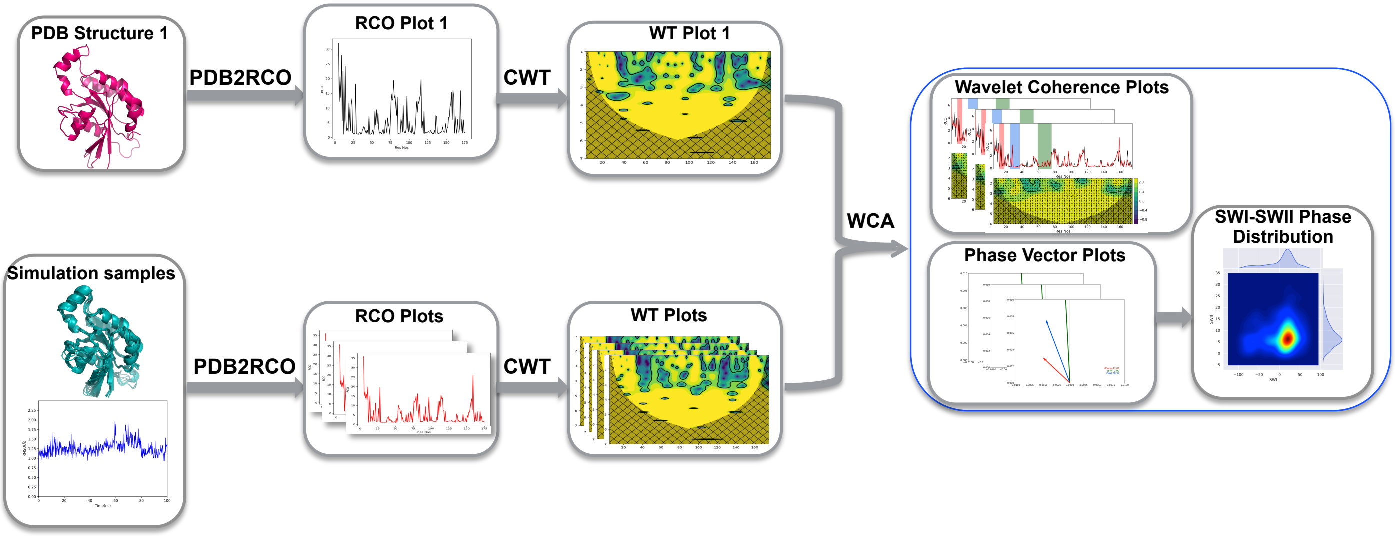

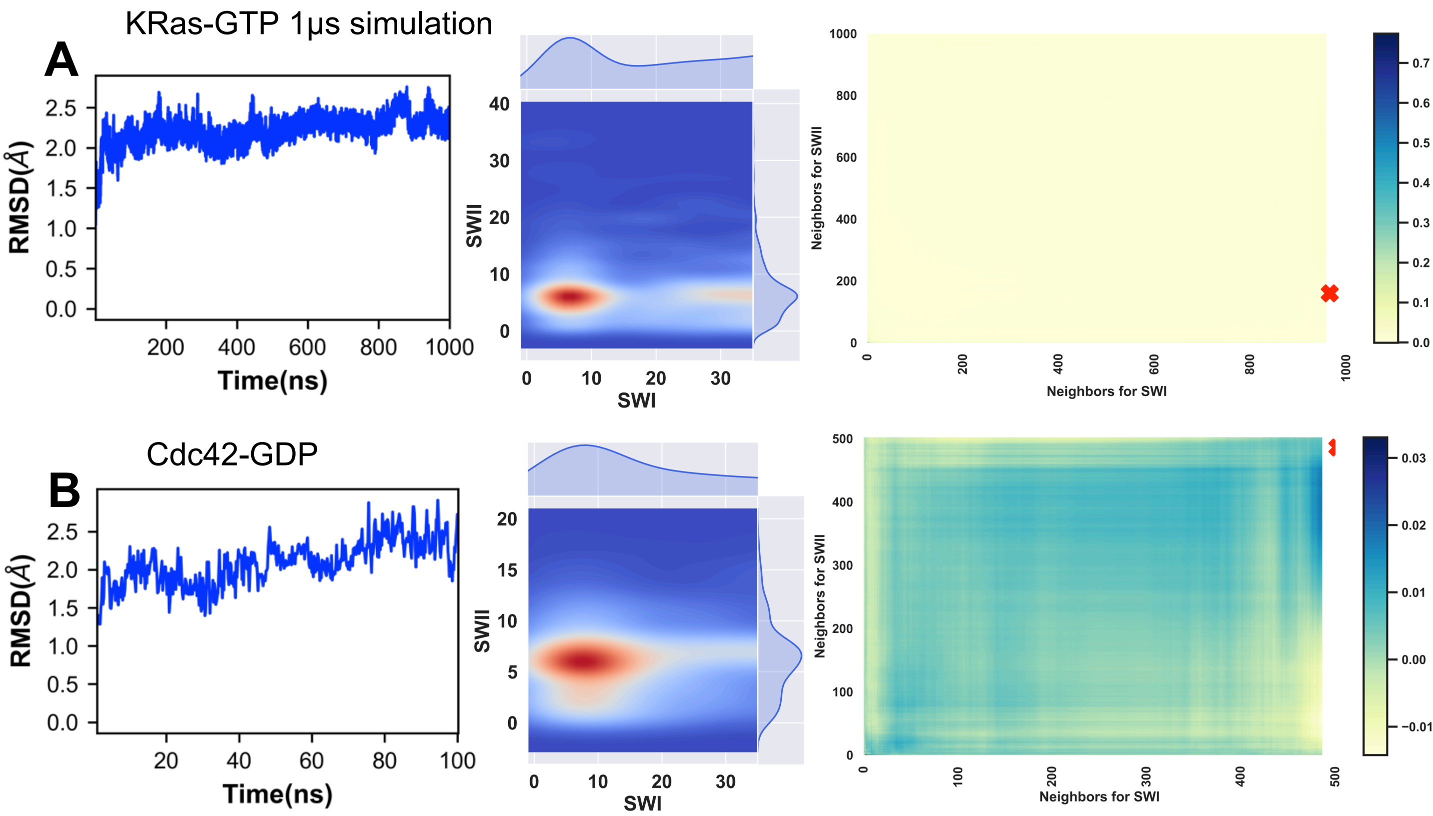
